# Home security cameras as a tool for behavior observations and science affordability

**DOI:** 10.1101/2023.04.17.537238

**Authors:** Billie C. Goolsby, Marie-Therese Fischer, Tony G. Chen, Daniela Pareja-Mejía, Daniel A. Shaykevich, Amaris R. Lewis, Gaelle Raboisson, Katharina Dellefont, Madison P. Lacey, Lauren A. O’Connell

**Author notes:** These authors contributed equally to the manuscript and mutually agree that authorship listing is interchangeable. To whom correspondence should be addressed: Billie C. Goolsby, Department of Biology, Stanford University, 371 Jane Stanford Way, Stanford, CA 94305, Lauren A. O’Connell, PhD., Department of Biology, Stanford University, 371 Jane Stanford Way, Stanford, CA 94305.

## Abstract

Reliably capturing transient animal behavior in the field and laboratory remains a logistical and financial challenge, especially for small ectotherms. Here, we present home camera systems as affordable, accessible, and suitable alternatives for monitoring small, cold-blooded animals historically overlooked by commercial camera traps. Home security cameras are often weather-resistant, operate offline or online, and allow collection of time-sensitive behavioral data in laboratory and field conditions with continuous data storage for up to four weeks. These lightweight cameras can also utilize phone notifications over Wi-Fi, alerting users when animals enter a space of interest and enabling sample collection at proper time periods. We present our findings, both technological and scientific, in an effort to elevate tools that enable researchers to maximize use of their research budgets. We discuss the relative affordability of our system for researchers in South America, home to the largest ectotherm diversity.

## Introduction

Accurately quantifying animal behavior is important for understanding how organisms respond to challenges and opportunities in their environment, which influences the fitness and survival of a species. Human presence can obscure natural behaviors (1,2) and remote cameras bypass the need for physical presence of the researcher. Remote camera approaches are used to study behavior at the population (3–6) and individual levels (7). Mammals and birds are the most common fauna monitored with cameras in the wild, but only a small range of prior studies (<2%) have focused on ectotherms, specifically amphibians and reptiles (8–12). This gap between ectotherms and endotherms can be explained by both technical and logistical difficulties, as many camera traps rely on body heat signatures (infrared, IR) and can be difficult to use in remote areas. Moreover, countries in the Global South hold much of the world’s ectotherm biodiversity, and yet accessibility and affordability of camera traps for local researchers can be prohibitive. For example, increasing camera weather resistance correlates with higher prices, and many camera traps also rely on cellular networks based in the Global North. An ideal camera system should be affordable, capable of detecting small ectotherm movement, weather-resistant, and operational in the absence of electricity and phone connectivity.

Here we describe multiple affordable camera systems originally designed for home security repurposed for research, which use pixel change rather than IR detection to trigger recording. We describe the specifications of these systems compared to three more commonly used camera traps in order for the reader to evaluate which recording device is best for their purposes. We benchmark our chosen camera set-up (both its trigger and 24/7 setting) against an established camera trap to highlight the suitability of home security cameras to monitor small ectotherms. We then present a case study on how this camera system can be used to quantify social behavior in small amphibians both in captivity and in the wild. Finally, we suggest efforts to make camera traps and recording devices more readily available and affordable for colleagues in the Global South as an example of how privileged institutions can actively contribute to making science more accessible.

## Results & Discussion

### Technical comparisons of remote monitoring systems for small ectotherms

We first compared technical specifications and availability of three classes of camera systems to evaluate the suitability for lab and field studies of amphibian behavior (**Table 1**). These include three monitoring systems (Reconyx [Holmen, WI, USA]; Bushnell [Overland Park, KS, USA], used in (9); GardePro [Hong Kong], used in (13)), three home security cameras (Wyze [Seattle, WA, USA], used herein and Google Nest [Mountain View, CA, USA], used in (14) and (15); and Tapo [Irvine, CA, USA]), and a manual camera system (Sony [Tokyo, Japan], used in (16)) To decide which two cameras to benchmark, our criteria focuses on ectotherm detectability, Wi-Fi availability, and affordability.

**Table 1.** Camera performance for studying ectotherm behavior. We compare features and accessibility of three home security systems (Wyze, Google Nest, Tapo) against four traditional camera traps for animal behavior currently on the United States commercial market (Sony, Reconyx Hyperfire 2, Bushnell Nature View HD, GardePro). If cameras are no longer commercially available by the original vendor, we provide the most similar commercially available camera from the same vendor. We discuss each feature in the context of studies on amphibian behavior. Abbreviations: LTE: long term evolution (a standard for wireless data transmission), HD: high definition, PIR: passive infrared sensor, IR: infrared, CMOS: Active pixel sensor; IP65 and IP66: indices for ingress protection (weather resistance, the higher the better), AA: single cell dry battery. Dark grey indicates home security cameras. Prices retrieved March 2026.

| System Specifications | Consequence on Amphibian Studies | Wyze Camera v3* | Google Nest Camera, Indoor & Wired | Tapo C120 | Sony Camera HDR-CX405 | Reconyx Hyperfire 2 Cellular Professional Covert IR Camera | Bushnell Nature View HD | GardePro A3S Trail Camera |
| --- | --- | --- | --- | --- | --- | --- | --- | --- |
| Ability to notify phone with local network | Quickly alerts researchers to behavior | Yes | Yes | Yes | No | Yes | No | No |
| Ability to notify phone <u>without</u> local network | Quickly alerts researchers to behavior | No | No | No | No | No | No | No |
| Cellular Source | Alerts rely on either local networks or cellular ones | 2.4 GHz local Wifi | 2.4GHz/5GHz) Wi-Fi | IEEE 802.11b/g/n, 2.4 GHz Wi-Fi | None | 3-4G LTE (ATT & Verizon, some international plans) | None | None |
| Price (USD) | Cost influences number of cameras purchasable | \$39.98 | \$99.99 | \$35.99 | \$229.99 | \$459.99 | \$299.99 | \$89.99 |
| Image Resolution | Equivalent across most cameras | HD 1920 x 1080p | HD (1920 x 1080) | 2K QHD 4MP (2560 x 1440 px) | 2.1 Megapixels (1920 x 1080) | 1080P HD Widescreen | HD 1080P | 48 Megapixels HD |
| Video Quality and sound | Equivalent across most cameras | HD 1920 x 1080p | HD (1920 x 1080) | 2K QHD 4MP (2560 x 1440 px, 20 fps) | 1920 x 1080/60p | 720P HD Video with Audio (cellular enabled cameras do not transmit video) | HD 1920x1080p | 2304 x 1296P@20fps (16:9 OR 1920 x |
|  |  |  |  |  |  |  |  | 1080P@30fps<br>(16:9)<br>OR<br>1280 x<br>720P@30fps<br>(16:9) |
| Detection Method<br>(Day) | Risk of not detecting<br>amphibians with PIR | Pixel change | Pixel change | 1/2.9"<br>Progressive<br>Scan CMOS<br>Starlight<br>Sensor | Exmor R<br>CMOS Sensor | PIR | PIR | PIR (high, normal<br>low) |
| Motion Detection<br>Video length | Variable, depends<br>on research goal | Up to 5 minutes<br>per notification | Up to 3 hours of<br>event history<br>totally | Up to 3<br>minutes per<br>notification | None, manual | Up to 90 seconds per<br>notification | up to 60s | Up to 5 minutes |
| Continuous Video<br>length | Remote monitoring<br>of behaviors that<br>may last longer than<br>several minutes. | 24/7 for up to 4<br>weeks with SD<br>card | 24/7 for up to 10<br>days with<br>subscription | 24/7 for up to 4<br>weeks with SD<br>card | Approximately<br>2 hrs 35<br>minutes of<br>battery life | No continuous<br>recording abilities<br>(trigger only) | No continuous<br>recording abilities<br>(trigger only) | No continuous<br>recording abilities<br>(trigger only) |
| Detection Method<br>(Night and<br>wavelength in nm) | IR wavelength that<br>can detect<br>amphibians | Near-IR (940)<br>and Far-IR (850) | Far-IR (850) | Near-IR (940)<br>and Far-IR<br>(850) | None explicitly<br>stated | No-Glow™ High<br>Output Covert IR | PIR | PIR (940nm) No<br>Glow IR LEDs |
| Camera weight (g) | Lower weights are<br>more carryable for<br>fieldwork | 98.8 | 393 | 100 | 190-215 | ~750 | 246.6 | 530 |
| IP Score | IP66 best for heavy<br>pressurized water<br>exposure, IP65<br>handles rain | IP65 | None provided. | IP66 | None provided. | IP66 | IP66 | IP66 |
| Operating<br>Temperatures (°C) | Most cameras<br>tolerate most biome temperature ranges | -20 - +40 | 0-+40 | -25 - +45 | None provided. | -40 - +60 | -4 - +140 | -20 - +60 |
| Night vision distance (m) | Depends on animal of interest | None provided | 15 | 30 | None provided. | 45 | 18.3 | 30 |
| Cloud saving abilities | Depends on researcher needs | Yes, with extra fees | Yes, with extra fees | Yes, with extra fees | No | No | No | No |
| Highest amount of internal memory allowed on camera via SD card (Gb) | Higher Gb allows for cameras being left alone longer | 512 | Up to 1 hr of internal storage (no Gb amount provided). | 512 | No limit for SD card | 512 | 32 | 512 |
| Power source (years last) | USB requires continuous power source, which needs to be resolved before field work | USB | USB | USB | Battery or USB | 12 AA (2 yrs) | AA (4-12)<br>Up to one year | AA (8)<br>Over a year depending on usage |
| Field of View | Depends on camera's ability to orient and point towards area of interest | 130° | 135° | 120°(Diagonal)<br>,<br>103°(Horizontal), 55°(Vertical) | 90-270° | None provided | 50° | Total 120° Central zone: 60°, Left side: 30°,Right side: 30° |

\* discontinued and replaced with Wyze v4 as of March 2026, both versions available on Amazon.

#### Ectotherm detection

The main challenge in remotely recording ectotherms is that many camera traps rely on passive infrared sensors (PIR) to detect thermal emissions (body heat) of endotherms (mainly mammals and birds) (9,17,18). Home security cameras frequently use pixel change, rather than IR detection, to trigger recording, which enables detection of ectotherms. Pixel change is broadly defined as a software’s comparison of pixel changes between photo or video frames, which is the basis for motion detection. In contrast to other systems we evaluated, both home security cameras Wyze v3 and Tapo has multiple degrees of night vision capabilities, including color night vision for dawn and dusk hours and for fully dark environments both near-infrared for close distance (940 nm) and far-infrared light range for far distance (850 nm). Home security cameras are not the only devices with pixel change detection methods, as the more traditional system Sony HDR-CX405 uses a CMOS sensor, albeit with a short battery life. In addition to PIR vs. pixel change ability, we also considered whether the camera could tolerate high-humidity and temperature conditions, since the majority of ectotherms live in tropical climates. Traditional cameras consistently outcompete home security cameras, frequently being IP66 (ingress protection index), which is considered the highest standard in weather resistance. Home security cameras that are designed for both indoor and outdoor use, although Tapo and Wyze are IP65, which are tolerant to tropical weather but would not be resistant to more extreme weather conditions like monsoons. Google Nest, which has no declared IP index, may be vulnerable to water damage.

#### Wi-Fi connection and power usage

An additional feature that is advantageous in the laboratory is the ability to signal to cellular devices to notify the user of a trigger. Many home security devices use the same technical infrastructure for monitoring filmed areas, such as a baby’s nursery or the front door to a home. Wyze v3, Google Nest, and Tapo connect to local networks rather than having 3G or 4G coverage, which is better for accessibility in countries in the Global South. Other considerations for remote recording of ectotherms are the device weight, power usage, and connectivity. Both Wyze and Tapo cameras are lightweight, which is critical for field applications where cameras are carried long distances to record for long periods of time. Notably, however, almost all home security cameras are wired, which present logistical challenges with fieldwork. This is not a challenge with traditional cameras. However, devices requiring power via USB can easily be co-packaged with battery packs for long-term fieldwork.

#### Affordability

The cost of cameras can be prohibitive, especially when multiple are needed at once for a field or laboratory study. Cameras can range from upwards of $450 USD per camera to $35 USD. Generally, home security cameras are around $50 USD per camera, whereas traditional camera traps have more price variation. Our study sites in the lab and in the field historically use a minimum of around four to five separate locations being separately monitored at the same time.

With these three criteria, we chose to compare the capabilities of GardePro and Wyze v3 within our laboratory, given that they represent traditional cameras and home camera systems, respectively. They were the most affordable of their categories at the time of our study while sharing similarities in motion detection length, storage capabilities, and weather resistance.

#### Benchmarking of Wyze v3 and GardePro

We sought to benchmark the abilities of a home security system (Wyze v3) to those of a more traditionally used camera, such as the GardePro A3S Trail Camera (GardePro, Tuen Mun, Hong Kong, **Fig 1**). Wyze has two different settings, continuous (24/7) to an SD card (secure digital card, holding up to 512 GB) and trigger, which responds to motion detection or sound only. We separately benchmarked with a recording period of 96 hours for each setting in a laboratory terraria that housed a large poison frog species, the Dyeing poison frog (*Dendrobates tinctorius*). We observed more behavior in Wyze continuous recordings than in GardePro (**Fig 1C**), although there was no significant difference between the seconds per observation by camera type (**Fig 1D**). On average, GardePro missed around 20.62 seconds of behavior compared to Wyze in continuous mode (**Fig 1E**). However, 24/7 recordings produce an abundance of data without behavior, which requires an experimenter to parse through.

**Fig 1.**
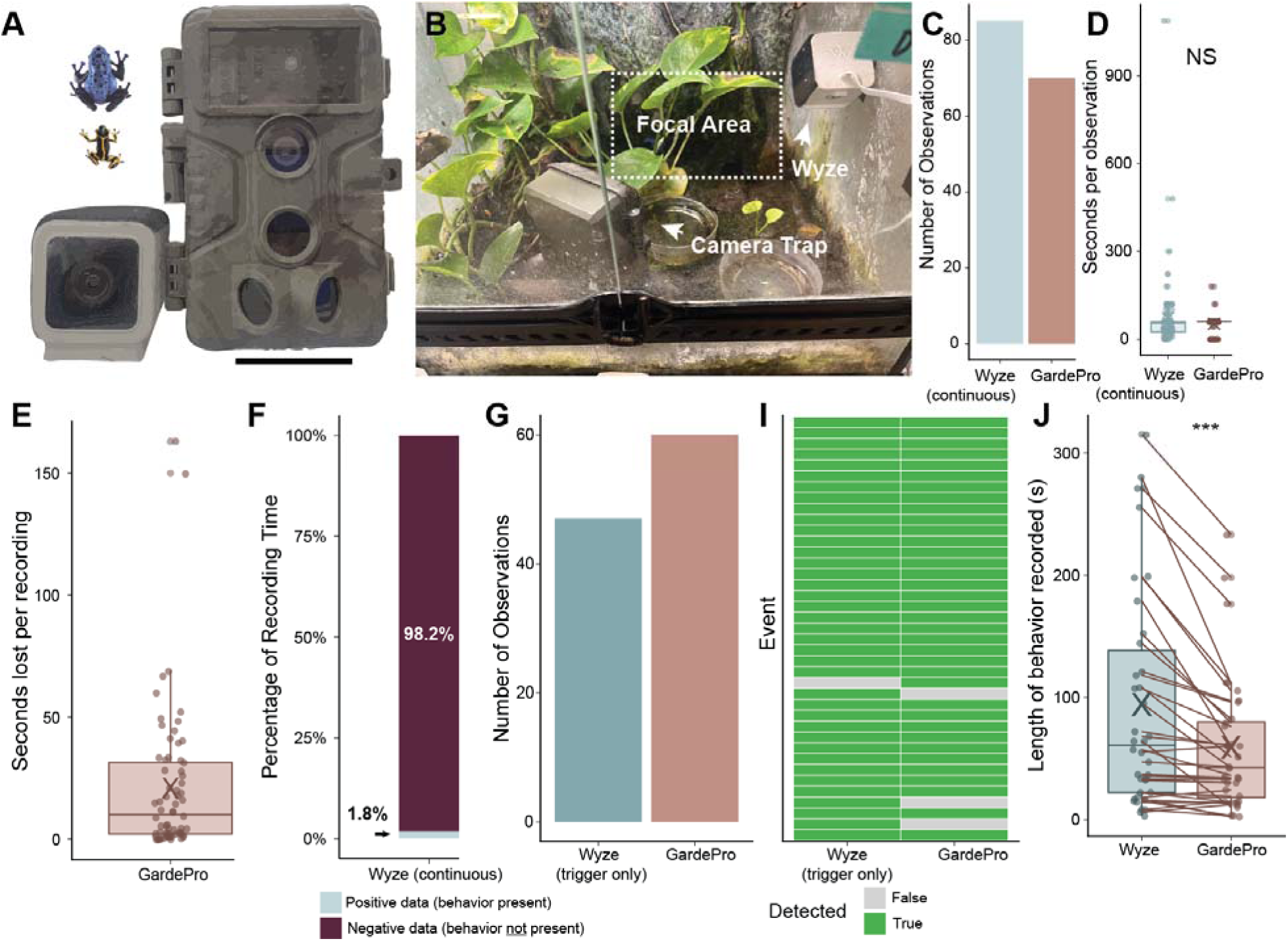
Comparisons of home security camera (Wyze) in both continuous (24/7) and trigger formats versus typical camera trap (GardePro) when recording the same area. **(A)** Size comparison of GardePro and Wyze v3 to two species of poison frogs, *Ranitomeya imitator* and *Dendrobates tinctorius.* Scale bar coincides to ∼52 mm. We **(B)** recorded *D. tinctorius* around a communal water pool and Coco-hut for a fairer comparison between both camera systems. **(C)** Number of observations detected by Wyze in continuous mode and GardePro. **(D)** Observation length in seconds per recording detected by Wyze in continuous mode and GardePro (no significant difference). **(E)** Due to Wyze recording in continuous mode, no data can be lost due to errors in triggering. However, 24/7 collection of data can produce an abundance of false negative data that takes a long time to parse through. On average, GardePro loses around 20 seconds of animal behavior per trigger compared to 24/7 recordings. We then compared this to Wyze’s continuous mode, which is constantly saving data. **(F)** In continuous mode, Wyze collects a disproportionately large amount of false positive data where no behavior is actually recorded (333900 seconds) compared to actual behavior (6153 seconds). **(G)** Wyze versus Gardepro’s trigger mechanisms. Wyze was triggered less often than GardePro, however, **(H)** GardePro fails to detect the same event that Wyze records slightly more often than Wyze fails to detect the same event that GardePro records. **(I)** Of the same events that GardePro and Wyze both successfully detected, Wyze recorded a significantly longer interval of behavior (p=2.54E-05,V=631, Wilcoxon signed rank test with continuity correction). To ensure equity in camera comparisons, we summed the amount of time GardePro or Wyze recorded total behavior per event, rather than per individual file, which would be due to storing mechanisms specific to each camera.

To more equitably compare cameras, we converted Wyze to trigger-only format (75% motion sensitivity). In this mode, we received less behavior triggers compared to GardePro (**Fig 1F**), although the Wyze camera more often detected the same behavior sequence (**Fig 1G**). To explain this discrepancy, we analyzed the recordings, finding that while Wyze was triggered less often, it recorded the detected behavior significantly longer (p=0.0003, t=3.97, **Fig 1H**). Following this benchmarking, we then sought to apply the system with the best overall performance to price ratio to quantify a more complex animal behavior within a laboratory environment.

### Proof of concept in the lab and field using Wyze v3

Given its size, affordability, and ectotherm detectability, we tested whether Wyze v3 can be used to record a complex social behavior under lab and field conditions. Specifically, we monitored the parental behavior of the variable poison frog (*Ranitomeya variabilis*) (**Fig 2**). In this species, eggs are laid terrestrially and then males transport their offspring to a pool of water, where the tadpole completes development (19). We used the Wyze camera system to record tadpole care and monitoring of available resources for tadpole deposition by adult frogs **(Fig 2A-C).** In field conditions, we were able to record the ecological context of this same behavior **(Fig 2D-E)**, which is important for knowing the ecological relevance of our laboratory findings. As field conditions differ markedly from laboratory environments (daily rainfall at the field site fluctuated between 0 to 30 mm and temperatures ranged from 20.5°C to 50°C), we systematically evaluated the cameras for user-friendliness, durability, quality of footage in terms of information gain, and their capacity to quantitatively capture behavior during offline operation. We further tested whether cameras could be broken apart readily to improve their focus on objects that are in proximity, given that home security cameras are designed to record at a focal point around 5-10 meters away. We were able to refocus cameras (**Fig 2F**, additional details at https://dx.doi.org/10.17504/protocols.io.dm6gp3p38vzp/v1) at no additional cost beyond a screwdriver.

**Fig 2.**
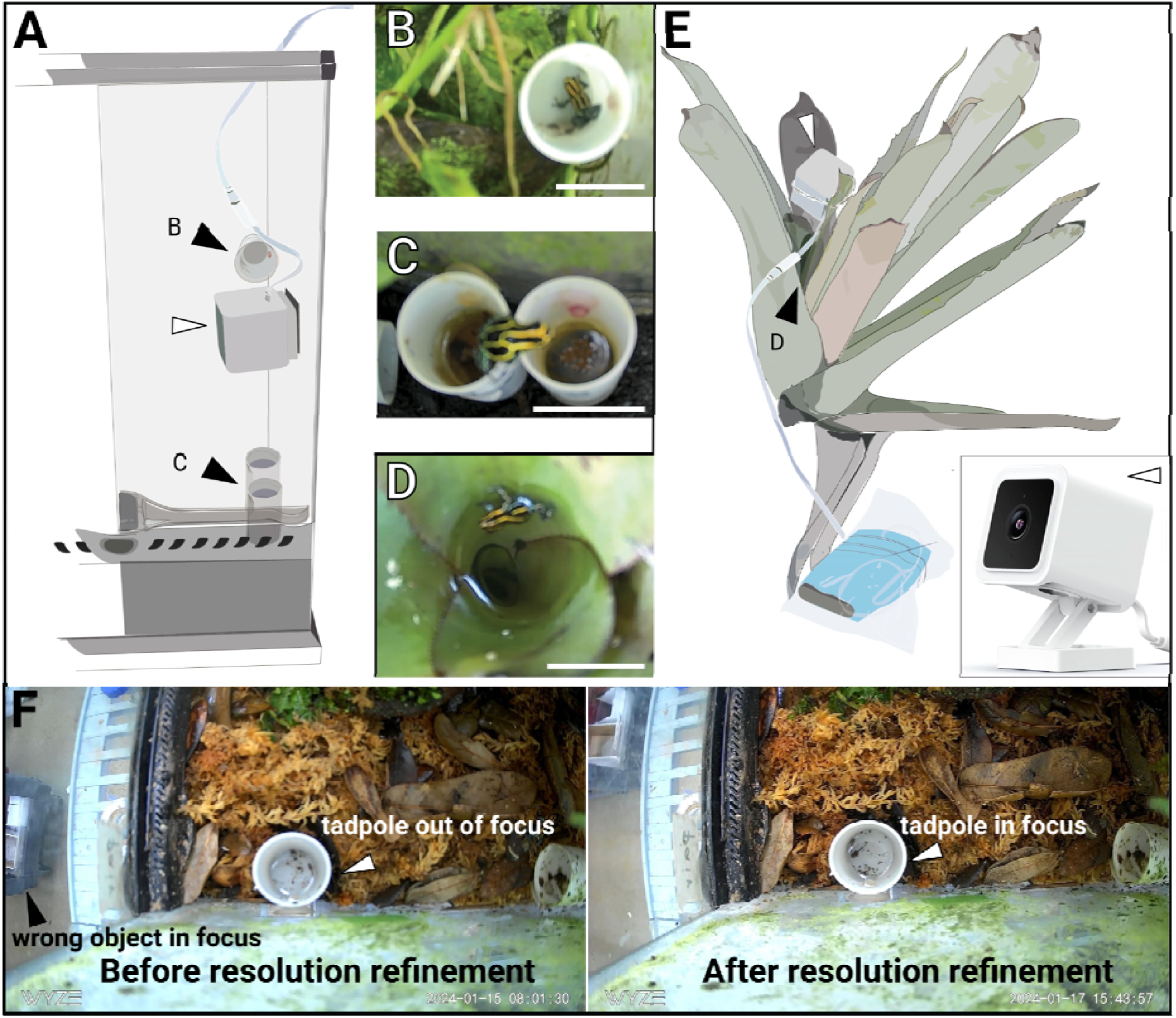
Camera installations in lab vivaria and in the field. **(A)** In lab vivaria, velcro was used to fi cameras above the desired focal subject, allowing for easy removal of the camera to switch SD cards. For monitoring parent-offspring interactions, the camera body (inlay, white arrow) was adjusted to focus on either the canisters containing the eggs that were mounted horizontally (inlay B) or the tadpole nursery (inlay C) (black arrows). The USB cables were connected to an outlet via a USB charging station. Inla pictures represent screenshots extracted from the recordings obtained from **(B)** egg attendance, **(C)** tadpole attendance in captivity and **(D)** tadpole attendance in the field. White scale bars represent 30 mm. **(E)** In nature, tadpoles grow up in water bodies formed by tightly overlapping leaf bases (black arrow, inlay (D) of large terrestrial plants called bromeliads. For observations, rubber bands were used to mount the cameras on bromeliad leaves above natural tadpole nurseries. The cameras (white arrows) were connected to the USB port of a power bank protected from the rain by two layers of plastic zip lock bags. Zoomed camera photo courtesy of Wyze Laboratories. **(F)** Cameras can be manually modified to improve resolution of focal sites, such as the tadpole inside of the plastic canister.

In the lab, we asked whether there were sex differences in parental attendance with egg clutches or tadpoles in two *R. variabilis* families (**Fig 3**). First, we monitored longitudinal parental behaviors from oviposition to tadpole transport (**Fig 3A,1B**). Only the father showed clutch attendance, where he visited daily, with a maximum of four times in a day, over a period of 10 days. He spent more time with his clutch for the first 3 days after deposition (roughly 10 min) compared to the last 3 days of his clutch’s development just before tadpole transport (roughly 5 min). These visits were typically spent touching or sitting on the eggs. In contrast, clutch visits by the female were limited to one brief event without physical contact with the clutch. During tadpole transport, the father transported each tadpole individually (n=3), with tadpole onboarding lasting 36 min on average. We also observed canister site fidelity for raising clutches, where the day after tadpole transport finished, the male resumed courtship and after two days of courting, the next clutch of eggs were laid.

**Fig 3.**
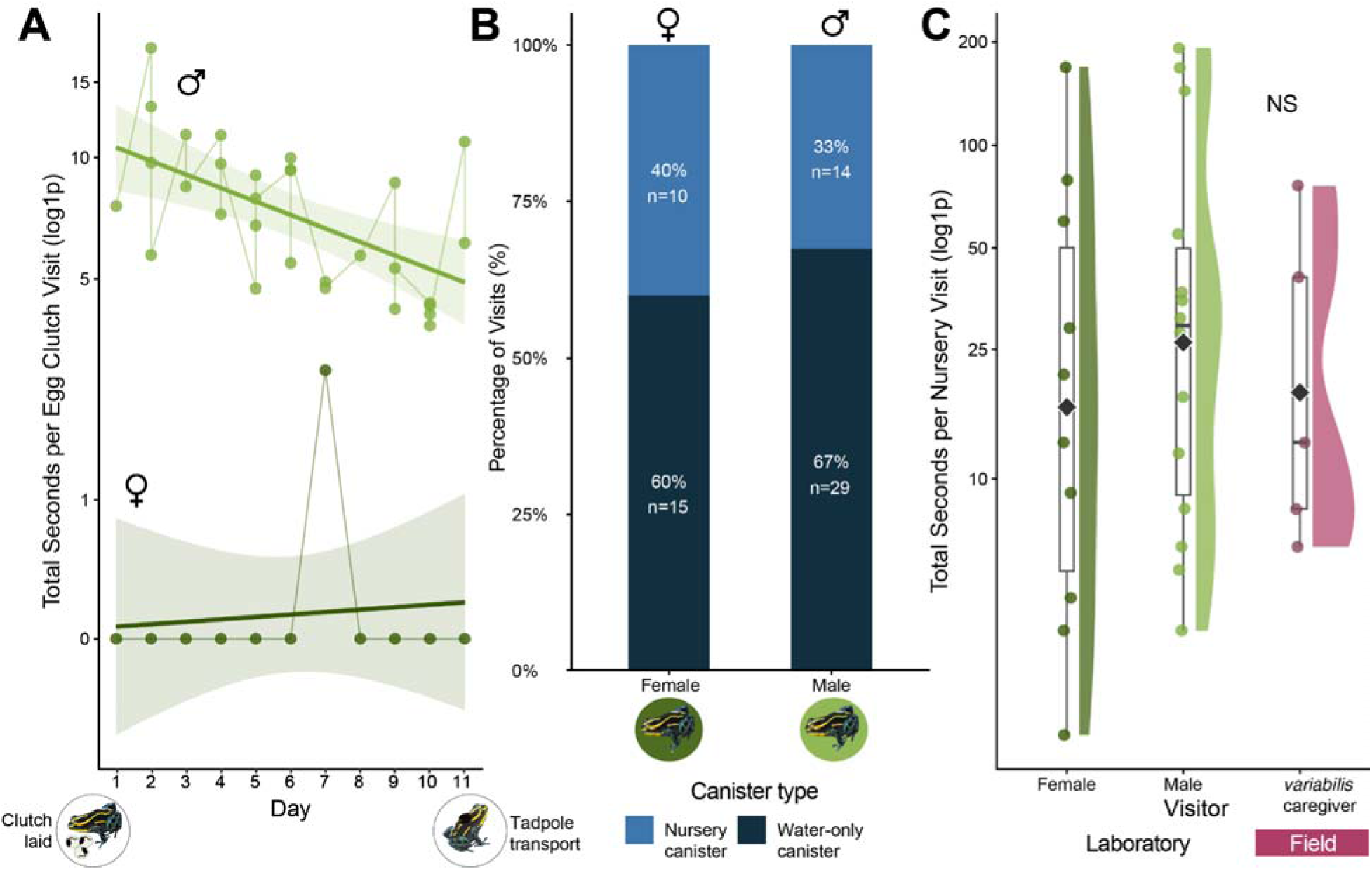
Comparisons of parental investment across offspring stage and in different environments. **(A)** Clutch attendance was male-only (light green), with visit duration highest immediately after egg deposition and declining as eggs matured. **(B)** Nursery visits (light blue) by male and female *R. variabilis* compared to visits to an adjacent unoccupied water canister (dark blue) during the same period. **(C)** Visit length did not differ significantly between captive males, captive females, or wild caregivers (purple; χ²=0.42, p=0.81, Kruskal-Wallis).

Next, we investigated the frequency and duration of parental visits to tadpole nurseries. Tadpole nurseries are small pools of water where poison frog fathers transport their tadpoles, which remain there until metamorphosis. To distinguish hydration and resource monitoring from parenting, we added an empty water pool adjacent to the nursery. We found that both the male and female preferred to visit unoccupied canister pools (**Fig 3B,1C**; 60% of female visits and 67% of male visits). Overall, the number and length of visits to laboratory tadpole pools were not significantly different in between male and female frogs (**Fig 3C**).

We then compared the frequency and length of nursery visits in the laboratory to those under natural conditions in the field. We monitored five tadpoles in bromeliad leaf axil pools for 10 days to compare parental behavior of *R. variabilis* in lab and field conditions. Although wild frogs visited tadpoles less frequently (5 visits) and for shorter durations (average of 29 seconds), these behavior traits did not differ significantly between frogs in the laboratory and those in the wild (χ^2^=0.42, p=0.81, Kruskal-Wallis rank sum test, **Fig 3C**). These findings indicate that parental care behavior in captive-bred frogs closely resembles that observed under natural conditions, supporting the comparability of behavioral observations from laboratory and field studies. Furthermore, these data provide the first evidence that *R. variabilis* regularly survey water bodies containing tadpoles in both lab and field conditions.

Image quality was sufficient to document parental behavior of *R. variabilis* under natural conditions and low-sensitivity operation mode was useful to record transient behaviors that are otherwise rarely observed, including predatory-prey interactions and territory disputes. These specifically included interactions with heterospecific intruders (**Supp Video 1**), predator encounters with spiders (**Supp Video 2**), aggressive behavior between tadpoles (**Supp Video 3**), and novel behaviors such as active wiping-off of transported offspring by adults (**Supp Video 4**), a behavior recently described in *Ameerega trivittata* (*20*) and not documented for *R. variabilis* until now, to our knowledge. Finally, we confirmed the camera could be used to record nocturnal ectotherm behavior by monitoring *R. variabilis* (**Fig 4C**).

**Fig 4.**
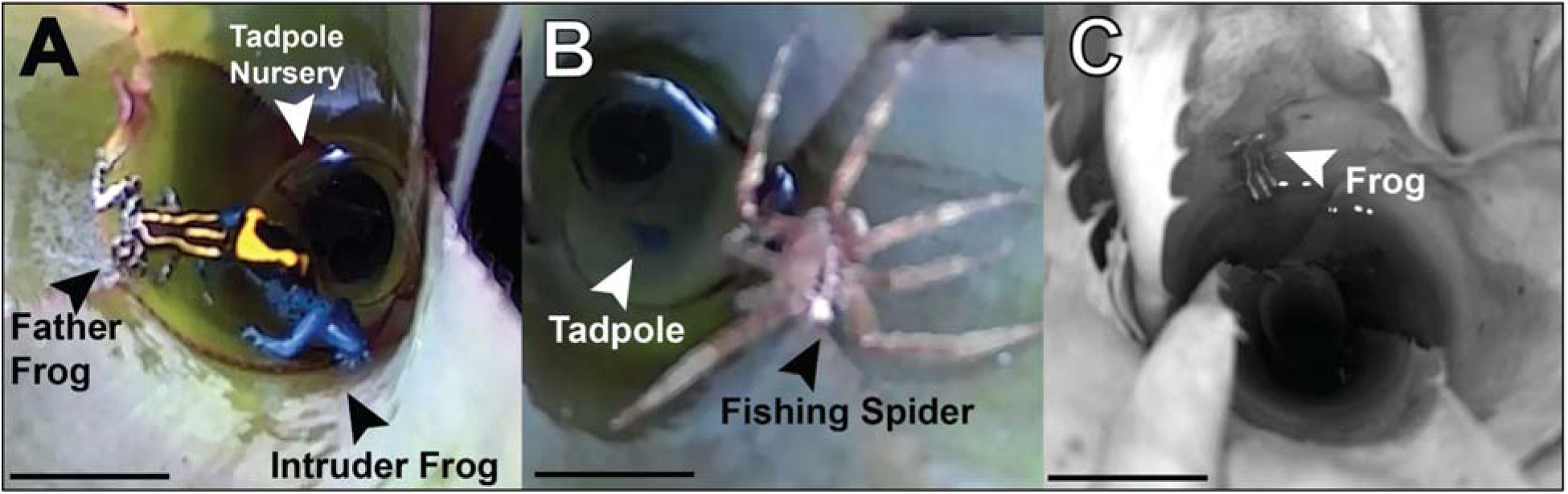
Remote observation captures of anuran behavioral ecology. **(A)** An *R. variabilis* male defends his tadpole nursery from a heterospecific intruder frog (*Dendrobates tinctorius*), a species with cannibalistic offspring. **(B)** An *R. variabilis* tadpole encounters a wandering spider (likely *genus Phoneutria*). **(C)** An *R. variabilis* investigates a bromeliad pool at night. (**A-C**) White and black scale bars represent 30 mm.

To further evaluate the applicability of home security cameras for monitoring arboreal amphibians, we deployed the Wyze v3 system to observe the behavior of the night-active fringed leaf frog, *Cruziohyla craspedopus*, in both field and laboratory conditions **(Fig 5**). In the laboratory, cameras were used to document general activity patterns rather than specific behavioral events. Individual frogs were recorded continuously within terraria for three consecutive days, allowing us to capture a broad range of behaviors such as climbing, resting, and exploratory movements throughout the enclosure (**Fig 5A**). As this species moves frequently across both vertical and horizontal axes in complex arboreal environments in the canopy, continuous tracking of individuals was very challenging in the field. Instead, cameras were positioned near potential reproductive sites, and branches frequently used by individuals, as well as at a focal tree where a resident frog was repeatedly observed (**Fig 5B**). This approach allowed opportunistic recording of natural behaviors without requiring continuous visual tracking of individuals. In several instances, cameras documented females approaching and interacting with these sites, as well as encounters between individuals occurring in proximity to egg-laying substrates. These observations allowed us to record egg clutches deposited on vegetation above water bodies, confirming that the monitored locations were actively used for reproduction. These recordings demonstrate that affordable security cameras can effectively document spontaneous activity and habitat use in highly mobile arboreal amphibians, even when precise focal tracking is not feasible.

**Fig 5.**
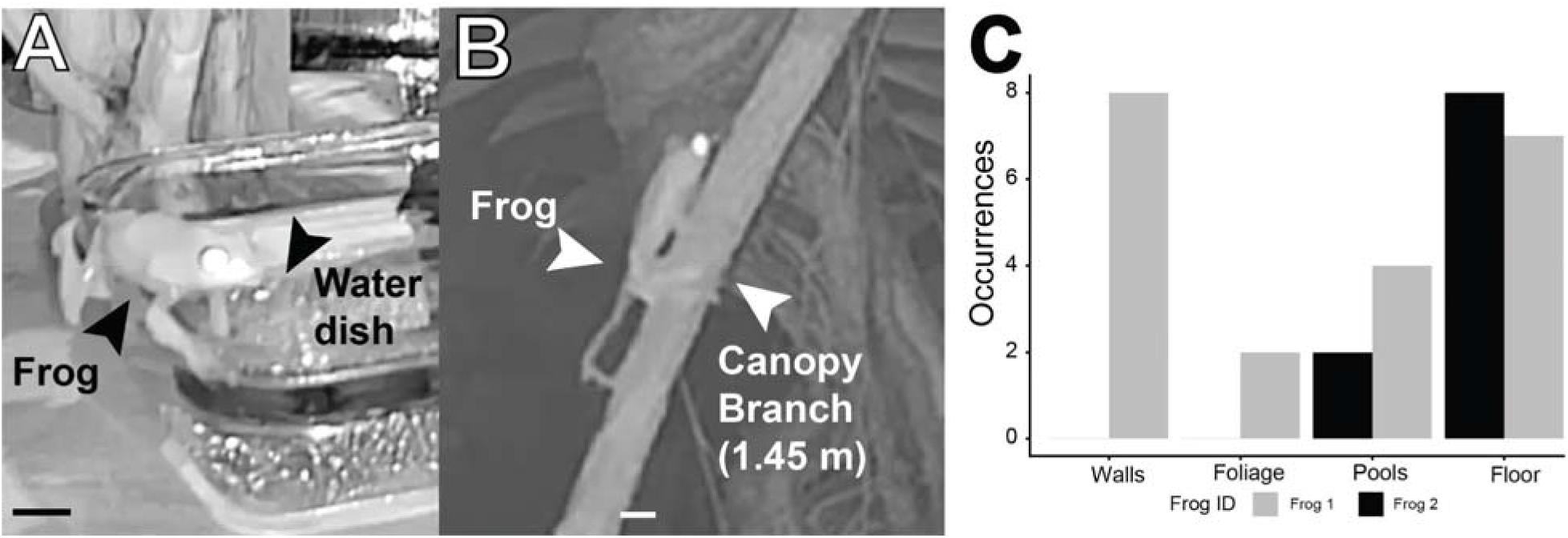
Observations of *C. craspedopus* nocturnal behavior. We recorded **(A)** nocturnal behavior of laboratory *C. craspedopus* colonies, such as attendance of water dishes. **(B)** Field observation of *C. craspedopus* frog climbing a canopy branch, with the frog and camera both at 1.45 m elevation. We **(C)** identified individual variation between laboratory frogs, such as two females (Frog 1 and Frog 2), where Frog 1 climbed walls and explored foliage more frequently, and Frog 2 was sedentary more often. No statistics were performed as this is intended to illustrate behavioral variation and usability for night recordings.**(A-B)** White and black scale bars represent 30 mm.

### Limitations of low-cost security cameras for field work

The cameras demonstrated good durability and handled exposure to extreme environmental conditions surprisingly well. Units deployed on exposed rock surfaces were additionally shielded from rain with simple plastic strips fixed above the openings without covering them. This simple adjustment was sufficient to prevent moisture ingress completely while not preventing rain from entering the tadpole nurseries. Three out of twenty units required replacement during a 1.5-month deployment: one camera cable was bitten by a small rodent, one began producing pink recordings after one month, and one yielded little footage despite no visible damage.

We learned many lessons when field testing camera systems, both positive outcomes and constraints. While Wi-Fi–dependent notifications are unavailable, the recording mode (continuous or motion-triggered) and the selected sensitivity level, once configured in the station with Wi-Fi, remained reliably preserved after disconnection and throughout offline use. The advantage of these cameras for behavioral monitoring lies in their affordability, and customizability, in particular, the ability to set low motion sensitivity to reduce data volume and analysis time. When sensitivity is set to high, light changes, moving vegetation and rainfall trigger recordings and generate tens of thousands of recordings per day; therefore, lowest sensitivity is required to maintain manageable datasets. The main practical limitation of the system was related to power cycling in the absence of Wi-Fi. Powering cameras with 26,800 mAh (96.48 Wh) power banks permitted 2 days of uninterrupted operation on both motion triggered and continuous mode. However, each time the camera was de- and reconnected to a power source in the field without Wi-Fi access, the internal timestamp reset to the factory default. This required manual correction of recording times by briefly filming a visible clock at startup and later using this recording to calculate the time offset. As each power bank replacement triggered a timestamp reset, the camera generated identical folder names at every restart, creating a risk of file overwriting. Thus, to prevent data loss, SD cards needed to be replaced every time the battery was changed, effectively doubling the number of SD cards required relative to the number of cameras. As recordings are automatically organized into multiple nested folders, managing large datasets required substantial time for sorting, renaming, and ensuring chronological accuracy across devices.

Given this limitation resulting from limiting camera sensitivity, we followed up on the reduced interaction frequency observed in the field by testing whether reduced numbers of visits reflected true behavior differences or resulted from missed events caused by the low-sensitivity setting. We positioned ten cameras above bromeliad leaf axils containing tadpoles, indicating that these axils are suitable sites for tadpole deposition. Inhabiting tadpoles were temporarily removed from their leaf axils and transferred to artificial canisters to empty preferred deposition sites for new deposits. Cameras were operated in motion-triggered mode at the lowest sensitivity setting. Batteries and SD cards were replaced every other day, and deposition events were independently verified by scoring the presence or absence of newly deposited small tadpoles each day. Of sixteen observed deposition events, six were captured on camera. This outcome suggests that while the low-sensitivity mode effectively reduces false triggers and minimizes data volume, it also fails to capture a substantial portion of behavioral events. As increasing the sensitivity rapidly led to an unmanageable number of recordings, we suggest using the cameras in continuous mode (potentially in combination with AI-assisted analysis) when quantitative data on behavior are required, and in low-sensitivity motion-triggered mode to obtain qualitative, complementary information on behavioral interactions without quantitative interpretation.

In summary, we found that affordable home security cameras can be on par or superior to other professional systems regarding image and video quality, detection methods, field of view and motion detection video length while being 3-18 times less expensive. Security cameras have been used to record behavior in frogs (21), fish (22), pigs (23), and chimpanzees (24). Recently, these cameras have also been paired with robotics to quickly and efficiently manipulate behavioral interactions when the camera was triggered (25). The diversity of animals and environments studied with these cameras illustrate their adaptability for many research projects.

### Further considerations for using camera systems for monitoring ectotherm behavior

While we found success in home security camera systems for our purposes and described their limitations, we highlight important further considerations for their use. First, in contrast to the Wyze v3, the newer Wyze v4 model was the only version readily available in many European countries. It offered improved image resolution and focus but was more expensive and retained the same limitations for offline use. In addition, full-length video recording and back-to-back event capture, both essential for fieldwork, required a subscription. Other added features, such as the built-in spotlight, were not useful for observing amphibian behavior. Second, our current study focused on one specific type of home security camera (Wyze v3), but multiple affordable alternatives can be possible, such as Reolink (New Castle, DE, USA) or Tapo/TP-Link (Irvine, CA). Additionally, recent efforts to make Raspberry Pi based-camera set ups allow for further customized camera trap creation (26). We found that the Tapo (**Table 1**) is as affordable as the Wyze, and also has a higher IP index, suggesting it may be more robust to weather conditions than Wyze and therefore useful across a wider range of experiments. Additionally, Tapo has a higher quality camera, which would make detection of ectotherms easier, making this option more viable than Wyze itself. We began our laboratory behavior studies before Tapo C120 was commercially available (early 2021 vs. late 2023/early 2024), which precluded our ability to test this option. Here, our approach of benchmarking camera performance under controlled laboratory conditions followed by experimental validation in the field provides a practical framework for evaluating the suitability of other camera systems for studies on ectotherm behavior.

### Camera resource affordability and access in scientific research

As ectotherms constitute more than 99% of species diversity (27,28), affordable remote camera systems would substantially improve approaches for studying behavior across a wide range of species. Much of the world’s ectothermic biodiversity is in the Global South, where research takes place within a context of historical and current colonialism and economic exploitation (29–31). Research is a fundamental activity taking place in almost every higher education institution, but differences in the level of science funding between the Global North and South impacts the types of questions that can be asked and how those questions will be answered (32). Affordable remote monitoring tools are especially critical in the Global South, where ectotherm biodiversity is greatest yet research infrastructure and equipment budgets are most constrained. While the Wyze v3/v4 is currently restricted to direct shipping within the United States and Canada, and import costs can approximately double the purchase price in countries such as Brazil, we present this system as a proof of concept that home security cameras as a class of device offer a viable, low-cost alternative to commercial camera traps. Equivalent pixel-change based systems are available through globally distributed retailers, including Reolink, Eufy, and Tapo/TP-Link.

An additional challenge to democratising home security camera systems for research includes language accessibility. To expand its usability, we also generated a Spanish translation of the Wyze v3 app and quick guides, available via GitHub and Protocols.io (<u>dx.doi.org/10.17504/protocols.io.dm6gp3p38vzp/v1</u>), to counteract the disadvantage incurred by the dominance of the English language in science (33). By creating these resources, we hope to make science more easily possible for all researchers. We suggest, therefore, that home security cameras like Wyze provide valuable, low-cost tools for documenting animal behavior in remote field conditions, but their application requires careful consideration of sensitivity settings, power logistics, and data management. Researchers in the Global North have an ethical responsibility to lead or participate in efforts that make scientific knowledge and tools more accessible through collaboration and shared resources.

## Methods

### Captive bred animals

All *Dendrobates tinctorius* used in the laboratory study were captive bred in our poison frog colony. One adult male and one adult female were housed together in a 45.72 x 45.72 x 45.72 cm terraria (Exoterra, Rolf C. Hagen USA, Mansfield, MA) containing sphagnum moss substrate, driftwood, live *Pothos* plants, thin petri dishes filled with water (treated with reverse osmosis R/O Rx, Josh’s Frogs, Owosso, MI) for tadpole deposition. Terraria were automatically misted ten times daily for 20 seconds each, and frogs were fed live *Drosophila hydeii* flies dusted with a vitamin powder and springtails three times per week. The observation housing was set on a 12:12 light cycle from 6:00 to 18:00. The average temperature and humidity of the observation was recorded for each day of observation, usually around 25°C and 95% humidity within the tank.

All *Ranitomeya variabilis* used in the laboratory study were captive bred in our poison frog colony or purchased from Ruffing’s Ranitomeya (Tiffon, Ohio, USA). One adult male and one adult female were housed together in a 45.72 x 30.48 x 30.48 cm terraria (Exoterra, Rolf C. Hagen USA, Mansfield, MA) containing sphagnum moss substrate, driftwood, live *Pothos* plants, horizontally mounted film canisters as egg deposition sites, and film canisters filled with water (treated with reverse osmosis R/O Rx, Josh’s Frogs, Owosso, MI) for tadpole deposition. Terraria were automatically misted ten times daily for 20 seconds each, and frogs were fed live *Drosophila melanogaster* flies dusted with a vitamin powder (Repashy Calcium Plus, and springtails three times per week. The observation housing was set on a 12:12 light cycle from 6:00 to 18:00. The average temperature and humidity of the observation was recorded for each day of observation, usually around 25°C and 95% humidity within the tank. Camera recordings of frogs in our captive breeding colony is approved by the Institutional Animal Care and Use Committee at Stanford University (protocols #34242).

### Field study site and wild study population

Cameras were tested for their usability in the field in a tropical primary rainforest in French Guiana. Testing was conducted in a natural study site situated on top of the mountain ‘Inselberg’ in vicinity to the Centre Nationale de la Recherche Scientifique managed research station (4°5’ N, 52°41’ W) within the Nature Reserve ‘Les Nouragueś. The study site is characterized by patches of Clusia trees separated by bare granite rocks and exposed to extreme environmental conditions (e.g. temperature oscillations between 18°C - 75°C) (34). The study site is inhabited by a population of *Ranitomeya variabilis* (CITES Appendix II, IUCN Conservation status: Least Concern). Camera recording of the study population on Inselberg was approved by the scientific committee of the Nouragues Ecological Research Station and under Stanford University Institutional Animal Care and Use Committee (protocol #33691).

*Ranitomeya variabilis* is a small dendrobatid poison frog that uses bromeliads as a resource for egg-laying and tadpole-rearing. Adult frogs are polygamous and typically lay clutches of 3-4 eggs into small arboreal bromeliads (*Catopsis berteroniana*) that are abundant on Clusia trees. Male frogs defend their territory and shuttle hatched tadpoles to individual water bodies that form in the leaf axils of the large terrestrial tank bromeliad (*Aechmea aquilega*). The cannibalistic tadpoles grow up in individual pools of 80 ml (35) where they feed on algae and leaf debris until they complete their development after about 3 months (36). While this species is not considered to engage in further parental care (19), frogs of this population have been documented to deposit fertilized eggs into bromeliad pools occupied by tadpoles at the start of the dry season (36). How frogs assess tadpole presence and resource availability to coordinate shuttling of offspring to unoccupied pools under these extreme environmental conditions is not known, presenting an ideal opportunity to test small camera traps in the field.

*Cruziohyla craspedopus*, the fringed leaf frog, is a visually striking amphibian native to the Amazonian rainforests of Colombia, Ecuador, Peru, and Brazil (37–39). This highly arboreal species spends the majority of its life in the forest canopy, where it forages for insects and avoids predators. Its fringed limbs and cryptic coloration facilitate camouflage within the complex canopy environment (40). Individuals descend to the forest floor only during the breeding season, seeking temporary pools or streams for reproduction (41). Individuals in the field were recorded in the Reserva Forestal Apayacu (1.06701° S, 77.65289° W) in Napo Province, Ecuador. Camera recording of *C. craspedopus* is approved under Stanford University Institutional Animal Care and Use Committee (protocol #34304).

### Camera setup and modifications

Detailed instructions on camera setup and usage can be found in the Supplementary materials on GitHub (https://github.com/laurenoconnelllab/WyzeSetUps).

In the laboratory, Wyze v3 cameras were adhered by velcro onto the side of the Exoterra tanks and suspended above the tadpole canister or focal area, with the face of the camera approximately 15.5-17.5 cm above the bottom of the canister or focal area. For egg care observations, Wyze v3 cameras are oriented horizontally, with the face of the camera facing approximately 10 cm from the canister entry site. Cameras were charged using their prepackaged USB cords and connected via a Nexwell USB charging station (Amazon, Bellevue, WA, USA) to the building outlets. Cameras were given 128 Gb SD cards for continuous recording. 128 Gb SD cards fill after approximately 10 days, and are switched by researchers and replaced with an empty SD card.

In the field, Wyze v3 cameras were equipped with a 128 GB SD card and set to the desired operating mode (motion triggered or continuous) using the WiFi of the field station before being transported to the field. We recommend setting the phone to the time zone in which the camera will be used during setup to avoid incorrect timestamps after changing time zones. Cameras were set to motion detection using the lowest sensibility, sound detection and notifications were switched off. All SD cards were formatted in the camera prior to use. Cameras were disconnected from power sources and WiFi during the transport to the study site. We used Anker PowerCore Fusion 10000 Model A1623 (9700mAh/35.21Wh) power banks for short-term (1 day) monitoring and Anker PowerCore26800 Model A1277 as well as 2nd Gen Astro E7 ModelA1210 (both 26800mAh/96.48Wh) for longer-term (10 days) surveil of tadpoles in their natural habitat. For long-term operation, we replaced the power bank every 2 days and recharged the devices in the camp. Prior to field implementation, we tested the efficiency of these power banks and found the operating mode of the camera (continuous vs motion-triggered) did not affect the monitoring time.

Cameras were labeled and secured to bromeliad leaves 20-40 cm above the water reservoir housing a tadpole (Fig 1E, black arrow) using commercial rubber bands (Fig 1E, white arrow). Power banks were wrapped with an absorbent diaper and placed in 2 plastic zip lock bags with their opening facing into different directions to minimize the amount of rain water seeping in (Fig 1E). We purposefully chose inexpensive zip lock plastic bags to cover the power supply from rain to test the operation of our system on a budget. Alternatively, plastic bags can be replaced by a variety of outdoor waterproof boxes designed for electrical equipment. We tried to shelter Power banks from the sun and rain by placing them under bromeliad leaves. To assure an accurate timestamp, we recorded the screen of our mobile phone showing the accurate time as a first action with the camera after reconnecting the camera to the power source. We replaced the SD card every 2 days when changing the power source to back up contained footage and moved the camera to a different tadpole nursery after 10 days.

As home security cameras are designed to film focal subjects at a distance (such as a driveway), filming animals within a close range presented a challenge. To improve resolution of objects at a closer distance, we dismantled the camera and shortened the lens distance. The protocol to replicate resolution modifications can be accessed on Protocols.io (<u>dx.doi.org/10.17504/protocols.io.dm6gp3p38vzp/v1</u>).

### Data Processing and Statistics

With continuous recording, every minute of every day is recorded. Videos were first processed and kept only when frogs entered the camera. The scorer marked the times the frog entered the focal canister, defining entrance and exit as when the entire frog’s body was within the canister. Data processing was performed in RStudio (version 4.2.3, PBC, Boston, MA). Kruskal-Wallis tests were performed for comparisons of frog attendance behavior, and Wilcoxon rank summed tests were performed for pairwise comparisons between GardePro and Wyze due to non-normal data distribution. Figures were created using the package ‘*ggplot2*’ (42) and in Adobe Illustrator (Adobe Illustrator 2023).

## Supporting information

Suppelmentary Video 2

## Acknowledgements

We gratefully acknowledge Dr. Michael Hobbs, who brainstormed with BG in the early stages of her PhD how to best detect our frogs. We thank the Wyze community forum, where members and the company together brainstorm new ideas and troubleshoot current technology. We thank David Ramirez for their care of our domestic poison frog colony. This research was conducted at Stanford University, which is located on the ancestral and unceded land of the Muwekma Ohlone tribe. Our field work was conducted in the Nature Reserve “Les Nouragues” in French Guiana, that was founded on land ancestrally inhabited by the Amerindien Nouragues (“Norak”) tribe. We thank the Nouragues research field station (managed by CNRS) which benefits from “Investissement d’Avenir” grants managed by Agence Nationale de la Recherche (AnaEE France ANR-11-INBS-0001; Labex CEBA ANR-10-LABX-25-01) and are extremely grateful to the camp managers and in particular Patrick Chatelet for his valuable technical, logistical and mental support throughout the field season.

## Data Availability

Field videos, raw data, and as well as code for running visualizations can be found in supplementary materials. Benchmarking videos are available on DataDryad. Detailed instructions for setting up cameras can be found on Github.

## Author Contributions

Conceptualization: BG, MTF, TGC

Data Curation: BG, MTF, GR, DPM, MPL

Formal Analysis: BG

Investigation: BG, MTF, GR, AL, DPM, TGC, DAS, MPL

Visualization: BG, MTF, TGC, DAS, DPM

Methodology: BG, MTF, DAS, TGC

Writing - Original Draft: BG, MTF, DPM

Writing - Review and Editing: BG, MTF, DPM, DAS, TGC, MPL, GR, AL, LAO

Project Administration: BG, MTF, LAO

Supervision: BG, MTF

Resources: LAO, MPL

Funding Acquisition: LAO

