## Supplementary Data and Code for "Home security cameras as a tool for behavior observations and science affordability": scriptforpaper_v2.R

```
#INSTALL NECESSARY PACKAGES 
library(ggplot2)
library(ggpubr)
library(ggrepel)
library(ggsci)
library(ggsignif)
library(plyr)
library(polynom)
library(pkgload)
library(pkgbuild)
library(pkgconfig)
library(purrr)
library(car)
library(corrplot)
library(gridExtra)
library(readxl)
library(scales)
library(glmmTMB)
library(DHARMa)
library(DescTools)
library(dunn.test)
library(tidyr)
library(dplyr)


#navigate to where "Recordings_Data.xlsx" is
setwd("/Users/billie/Downloads")

#Report data from the laboratory observations, combined across both egg canisters and tadpole pools##
CombinedVisits <- read_excel("Recordings_Data.xlsx", sheet = "CombinedVisits")

#Report data from the laboratory observations of counting how many visits occured across empty canisters and canisters filled with water#
tadpolevisitcounts <- read_excel("Recordings_Data.xlsx", sheet = "TadpolePoolCounts")

# Reshape to long format

tadpolevisitcounts_long <- tadpolevisitcounts %>%
  pivot_longer(cols = c(FemalePercentage, MalePercentage),
               names_to = "sex",
               values_to = "percentage") %>%
  mutate(sex = recode(sex,
                      "FemalePercentage" = "Female",
                      "MalePercentage"   = "Male"))

# Combined stacked bar
tad_combined <- ggplot(tadpolevisitcounts_long, 
                       aes(x = sex, y = percentage, fill = `...1`)) +
  geom_bar(stat = "identity", width = 0.5) +
  geom_text(aes(label = percent(percentage / 100)),
            position = position_stack(vjust = 0.5),
            size = 3, color = "white", fontface = "bold") +
  scale_fill_manual(values = c("#3E76AD", "#234261")) +
  theme_classic() +
  theme(text = element_text(size = 12),
        legend.position = "bottom") +
  labs(x = NULL, y = "Percentage of Visits (%)", fill = NULL)

tad_combined <- ggplot(tadpolevisitcounts_long,
                       aes(x = sex, y = percentage, fill = `...1`)) +
  geom_col(width = 0.5) +
  geom_text(aes(label = paste0(round(percentage), "%")),  # fix: was dividing already-decimal value
            position = position_stack(vjust = 0.5),
            size = 3.5, color = "white", fontface = "bold") +
  scale_fill_manual(values = c("Tadpole" = "#3E76AD", "Water" = "#1E4D7A"),
                    labels = c("Tadpole", "Water")) +
  scale_y_continuous(labels = scales::percent_format(scale = 1)) +
  theme_classic() +
  theme(text = element_text(size = 12),
        legend.position = "bottom") +
  labs(x = "Visitor Sex", y = "Percentage of Visits (%)", fill = NULL)  # fix: "Visitor Sex" not blank

tad_combined

###Comparing how the sexes visit tadpoles in the amount of time invested, and comparign this with our field data###


tadpolepoolvisits_tadpoleonly <- read_excel("Recordings_Data.xlsx",sheet = "TadpolePoolVisits_tadpoleonly", col_types = c("date", "date", "date", "date", "text", "text", "text", "numeric","text", "text","text"))
tadpolepoolvisits_tadpoleonly$visitorsex<- as.character(tadpolepoolvisits_tadpoleonly$visitorsex)
tadpolepoolvisits_tadpoleonly$visitorsex<- factor(tadpolepoolvisits_tadpoleonly$visitorsex,levels=unique(tadpolepoolvisits_tadpoleonly$visitorsex))

vplotlab <- ggplot(tadpolepoolvisits_tadpoleonly, aes(x=`visitorsex`, y=`Totalsecondsduration`, fill = `visitorsex`, color = `visitorsex`))  +
  scale_fill_manual(values = c("#365902","#81AE3E","#AD3E66"))  + 
  scale_color_manual(values = c("#5A9403","#365902","#83455C")) +
  geom_boxplot() + 
  stat_summary(fun=mean, geom="text",label="X",size=6)+ 
  geom_jitter(size=3, position= position_jitterdodge()) + 
  theme_classic() +
  theme(text = element_text(size = 13), panel.border = element_blank(), panel.grid.major = element_blank(),panel.grid.minor = element_blank())+ labs(x = "Sex", y = "Total Seconds per Nursery Visit")
vplotlab

library(ggdist)

vplotlab_rain <- ggplot(tadpolepoolvisits_tadpoleonly, 
                        aes(x = visitorsex, y = Totalsecondsduration, 
                            fill = visitorsex, color = visitorsex)) +
  
  # Half violin (the "cloud")
  stat_halfeye(adjust = 0.5, width = 0.4, justification = -0.3,
               .width = 0, point_colour = NA, alpha = 0.7) +
  
  # Boxplot (thin, minimal)
  geom_boxplot(width = 0.12, outlier.shape = NA, alpha = 0.4,
               color = "grey30", fill = "white") +
  
  # Raw points (the "rain")
  geom_jitter(width = 0.05, size = 2.5, alpha = 0.8) +
  
  # Mean as a diamond
  stat_summary(fun = mean, geom = "point", shape = 23,
               size = 4, color = "white", fill = "grey20") +
  
  scale_fill_manual(values  = c("#365902", "#81AE3E", "#AD3E66")) +
  scale_color_manual(values = c("#365902", "#81AE3E", "#83455C")) +
  
  theme_classic() +
  theme(
    text = element_text(size = 13),
    legend.position = "none",          # sex is already on x-axis
    axis.line = element_line(colour = "black"),
    panel.grid = element_blank()
  ) +
  labs(x = "Sex", y = "Total Seconds per Nursery Visit")

vplotlab_rain


vplotlab_rain <- ggplot(tadpolepoolvisits_tadpoleonly,
                        aes(x = visitorsex, y = Totalsecondsduration,
                            fill = visitorsex, color = visitorsex)) +
  stat_halfeye(adjust = 0.5, width = 0.4, justification = -0.3,
               .width = 0, point_colour = NA, alpha = 0.7) +
  geom_boxplot(width = 0.12, outlier.shape = NA, alpha = 0.4,
               color = "grey30", fill = "white") +
  geom_jitter(width = 0.05, size = 2.5, alpha = 0.8) +
  stat_summary(fun = mean, geom = "point", shape = 23,
               size = 4, color = "white", fill = "grey20") +
  scale_x_discrete(labels = c("female" = "Female",         # fix: clean up capitalisation
                              "male"   = "Male",
                              "fieldfrog" = "Field Frog")) +
  scale_y_continuous(trans = "log1p",                      # fix: compresses outliers at 170-190s
                     breaks = c(0, 10, 25, 50, 100, 200),
                     labels = c(0, 10, 25, 50, 100, 200)) +
  scale_fill_manual(values  = c("#365902", "#81AE3E", "#AD3E66")) +
  scale_color_manual(values = c("#365902", "#81AE3E", "#83455C")) +
  theme_classic() +
  theme(text = element_text(size = 13),
        legend.position = "none") +
  labs(x = "Visitor Sex",                                  # fix: was "Sex", fieldfrog is still a sex category
       y = "Total Seconds per Nursery Visit (log1p)")

kruskal.test(Totalsecondsduration ~ visitorsex, data = tadpolepoolvisits_tadpoleonly)
dunn.test(tadpolepoolvisits_tadpoleonly$Totalsecondsduration,
          tadpolepoolvisits_tadpoleonly$visitorsex,
          method = "bonferroni",   # correction method
          kw = TRUE,               # prints KW result again for reference
          label = TRUE,            # labels output with group names
          wrap = FALSE)
###Comparing how the sexes visit egg canisters during development until tadpole transport, which happens on day 12###


Combinedvisitseggsonly_lessthan <- read_excel("Recordings_Data.xlsx", sheet = "CombinedVisitsEggsOnly_lessthan")
#CombinedVisits_eggsonly$date.factor <- factor(CombinedVisits_eggsonly$date)
#CombinedVisits_eggsonly$day <- factor(CombinedVisits_eggsonly$day)

#CombinedVisits_eggsonly$day <- as.Date(CombinedVisits_eggsonly$day, format = '%d')
vplotlab2 <- ggplot(Combinedvisitseggsonly_lessthan, aes(x=as.numeric(`day`), y= as.numeric(`NumberofMinutes`), group = `visitorsex`, fill = `visitorsex`, color = `visitorsex`)) +
  scale_fill_manual(values = c("#365902","#81AE3E","#CB4978"))  + 
  scale_color_manual(values = c("#365902","#81AE3E","#83455C")) +
  geom_line() + 
  geom_point() + 
  geom_smooth(method = "lm") + 
  theme_classic() + theme(text = element_text(size = 13), panel.border = element_blank(), panel.grid.major = element_blank(),panel.grid.minor = element_blank()) + 
  labs(x = "Sex", y = "Total Seconds per Nursery Visit")
vplotlab2

vplotlab2 <- ggplot(Combinedvisitseggsonly_lessthan,
                    aes(x = as.numeric(day),
                        y = as.numeric(NumberofMinutes),
                        fill = visitorsex,
                        color = visitorsex)) +
  geom_smooth(method = "lm", alpha = 0.15, linewidth = 1.2) +
  geom_line(alpha = 0.4) +
  geom_point(size = 2.5, alpha = 0.8) +
  scale_y_continuous(trans = "log1p",                     # fix: handles 0s and outlier compression
                     breaks = c(0, 1, 5, 10, 15),
                     labels = c(0, 1, 5, 10, 15)) +
  scale_fill_manual(values  = c("#365902", "#81AE3E", "#CB4978")) +
  scale_color_manual(values = c("#365902", "#81AE3E", "#83455C")) +
  theme_classic() +
  theme(text = element_text(size = 13),
        strip.background = element_blank(),
        strip.text = element_text(face = "bold"),
        legend.position = "none",
        panel.spacing = unit(1, "lines")) +
  labs(x = "Day", y = "Total Seconds per Egg Clutch Visit (log1p)")  # fix: was "Sex"


grid.arrange(tad_combined,vplotlab2,vplotlab_rain,nrow=1)

######wyze positive vs negative. results######
continuous_wyze <- read_excel("Perch_comparisons.xlsx",  sheet = "contiinuous_wyze")

continuous_wyze$percentage <- continuous_wyze$seconds / sum(continuous_wyze$seconds) * 100

continuouspositivevsnegative <-ggplot(continuous_wyze, aes(x = camera, y = percentage, fill = category)) +
  geom_col(width = 0.4) +
  geom_text(aes(label = paste0(round(percentage, 1), "%")),
            position = position_stack(vjust = 0.5),
            color = "white", fontface = "bold", size = 4) +
  scale_fill_manual(values = c("positive" = "#C1D6D9", "negative" = "#59253E")) +
  scale_y_continuous(labels = scales::percent_format(scale = 1)) +
  theme_classic(base_size = 13) +
  labs(x = NULL, y = "Percentage of Recording Time",
       fill = "Detection") + theme(legend.position = "none")


######wyze vs gardpro comparisons######

perch_comparisons_continuous <- read_excel("Perch_comparisons.xlsx",  sheet = "continuous")
perch_comparisons_continuous$secondsoff[trimws(perch_comparisons_continuous$secondsoff) == ""] <- NA
perch_comparisons_continuous$camera <- 
  factor(perch_comparisons_continuous$camera,
         levels = c("247wyze", "gardepro"))


perch_comparisons_continuous_gardepro <- subset(perch_comparisons_continuous, camera=='gardepro')

mean(perch_comparisons_continuous_gardepro$secondsoff, na.rm = TRUE)

continuous_secondslost <- ggplot(data=perch_comparisons_continuous_gardepro, aes(x=camera, y=secondsoff, fill = camera, color = camera)) +
  scale_fill_manual(values = c("#BF8F84","#C1D6D9"))  + 
  scale_color_manual(values = c("#734E46","#80A2A6")) +
  geom_boxplot(alpha = 0.5) +
  ylim(-1,200)+
  geom_jitter(width = 0.1, size = 1.5, alpha = 0.6) +
  stat_summary(fun=mean, geom="text",label="X", size=6)+ 
  theme_classic() + theme(text = element_text(size = 13), panel.border = element_blank(), panel.grid.major = element_blank(),panel.grid.minor = element_blank()) + 
  labs(y = "Seconds lost per recording") + theme(legend.position = "none")

continuous_secondslost

continuous_timedifference <- ggplot(perch_comparisons_continuous,
                                    aes(x = camera, y = timedifferenceinseconds,
                                        fill = camera, color = camera)) +
  scale_fill_manual(values = c("#C1D6D9","#BF8F84")) +
  scale_color_manual(values = c("#80A2A6","#734E46")) +
  geom_boxplot(alpha = 0.5) +
  geom_jitter(width = 0.1, size = 1.5, alpha = 0.6) +
  stat_summary(fun=mean, geom="text",label="X", size=6) +
  theme_classic() +
  labs(y = "Seconds lost per recording") +
  theme(text = element_text(size = 13),
        legend.position = "none")

continuous_timedifference


continuous_observations_counts <- ggplot(perch_comparisons_continuous, aes(x = camera, fill = camera)) +
  geom_bar() +
  scale_fill_manual(values = c("#C1D6D9","#BF8F84")) +
  theme_classic() +
  labs(x = "Camera", y = "Count") + theme(legend.position = "none")

continuous_observations_counts

perch_comparisons_trigger <- read_excel("Perch_comparisons.xlsx",  sheet = "trigger")

perch_comparisons_trigger$camera <- 
  factor(perch_comparisons_trigger$camera,
         levels = c("wyze", "gardepro"))

triggers_observations <- ggplot(perch_comparisons_trigger, aes(x = camera, fill = camera)) +
  geom_bar() +
  scale_fill_manual(values = c("#BF8F84","#82A8AE")) +
  theme_classic() +
  labs(x = "Camera", y = "Count")+ theme(legend.position = "none")

triggers_observations

grid.arrange(continuous_observations_counts, continuous_timedifference, continuous_secondslost, triggers_observations, nrow= 1)

####

ggplot(data=perch_comparisons_trigger, aes(x=camera, y=timedifferenceinseconds, fill = camera, color = camera)) +
  scale_fill_manual(values = c("#82A8AE","#BF8F84"))  + 
  scale_color_manual(values = c("#3F5558","#734E46")) +
  geom_boxplot() + geom_jitter() +
  stat_summary(fun=mean, geom="text",label="X",size=6)+ 
  theme_classic() + theme(text = element_text(size = 13), panel.border = element_blank(), panel.grid.major = element_blank(),panel.grid.minor = element_blank()) + 
  labs(x = "Camera Type", y = "Length of Individual Recordings (s)")


perch_comparisons_trigger_summedup <- read_excel("Perch_comparisons.xlsx",  sheet = "trigger_summedup")

perch_comparisons_trigger_summedup_plot <- ggplot(data=perch_comparisons_trigger_summedup, aes(x=Camera, y=TimeInSeconds, fill = Camera, color = Camera)) +
  scale_fill_manual(values = c("#BF8F84","#82A8AE"))  + 
  scale_color_manual(values = c("#734E46","#3F5558")) +
  geom_boxplot(alpha = 0.5) +
  geom_jitter(width = 0.1, size = 1.5, alpha = 0.6) +
  geom_line(aes(group= EventPair)) +
  stat_summary(fun=mean, geom="text",label="X",size=7)+ 
  theme_classic() + theme(text = element_text(size = 13), panel.border = element_blank(), panel.grid.major = element_blank(),panel.grid.minor = element_blank()) + 
  labs(x = "Camera Type", y = "Length of behavior recorded (s)") + theme(legend.position = "none")

perch_comparisons_trigger_summedup_plot

t.test(TimeInSeconds ~ Camera,
       data = perch_comparisons_trigger_summedup)

grid.arrange(continuous_observations_counts, continuous_timedifference, continuous_secondslost, triggers_observations, perch_comparisons_trigger_summedup_plot, nrow= 2)

library(tidyr)
library(dplyr)

df_wide <- perch_comparisons_trigger_summedup %>%
  group_by(EventPair, Camera) %>%
  summarize(TimeInSeconds = mean(TimeInSeconds), .groups = "drop") %>%
  pivot_wider(names_from = Camera,
              values_from = TimeInSeconds,
              values_fill = 0)

t.test (df_wide$wyze,
       df_wide$gardepro,
       paired = TRUE)

wilcox.test(df_wide$wyze, df_wide$gardepro, paired = TRUE)
  
perch_comparisons_trigger_summedup %>%
  count(EventPair) %>%
  table()

df_plot <- perch_comparisons_trigger_summedup %>%
  group_by(EventPair, Camera) %>%
  summarize(TimeInSeconds = mean(TimeInSeconds), .groups = "drop") %>%
  complete(EventPair, Camera, fill = list(TimeInSeconds = 0)) %>%
  mutate(detected = TimeInSeconds > 0)

detection_plot <- ggplot(df_plot, aes(x = Camera, y = factor(EventPair), fill = detected)) +
  geom_tile(color = "white") +
  scale_fill_manual(values = c("TRUE" = "#4CAF50", "FALSE" = "#D3D3D3")) +
  labs(x = "Camera", y = "Event", fill = "Detected") +
  theme_classic() +
  theme(legend.position="none")+
  theme(axis.text.y = element_blank(),
        axis.ticks.y = element_blank())

detection_plot

df_plot %>%
  group_by(Camera) %>%
  summarize(detection_rate = mean(detected)) %>%
  ggplot(aes(x = Camera, y = detection_rate, fill = Camera)) +
  geom_col() +
  ylim(0,1) +
  labs(y = "Detection rate") +
  theme_classic() +
  theme(legend.position = "none")

grid.arrange(continuous_observations_counts, continuous_timedifference, continuous_secondslost, continuouspositivevsnegative, triggers_observations, detection_plot, perch_comparisons_trigger_summedup_plot, nrow= 2)

tight <- theme(
  plot.margin = unit(c(0.1, 0.1, 0.1, 0.1), "cm")
)

p1 <- continuous_observations_counts + tight
p2 <- continuous_timedifference + tight
p3 <- continuous_secondslost + tight
p4 <- continuouspositivevsnegative + tight
p5 <- triggers_observations + tight
p6 <- detection_plot + tight
p7 <- perch_comparisons_trigger_summedup_plot + tight

grid.arrange(p3,p4,p5,p6,p7, nrow=1)

grid.arrange(
  p1, p2, p3, p4,
  p5, p6, p7,
  layout_matrix = layout,
  heights = c(1, 1),
  widths  = c(1, 1, 1, 1)
)


Cruziotable <- read_excel("Perch_comparisons.xlsx",  sheet = "table")

df <- data.frame(
  Frog = c("CC006", "CC011"),
  `Use Floor` = c(7, 8),
  `Use Walls` = c(8, 0),
  `Use Plants` = c(4, 2),
  `Use Pool` = c(2, 0),
  check.names = FALSE
)

df_long <- df %>%
  pivot_longer(-Frog, names_to = "Behavior", values_to = "Count")

ggplot(df_long, aes(x = Behavior, y = Count, fill = Frog)) +
  geom_bar(stat = "identity", position = "dodge") +
  scale_fill_manual(values = c("CC006" = "grey", "CC011" = "black")) +
  labs(x = "Behavior", y = "Occurrences", fill = "Frog ID") + 
  theme_classic()+theme(legend.position = "bottom")
```
